# Control of testes mass by androgen receptor paralogs in a cichlid

**DOI:** 10.1101/2021.02.23.432504

**Authors:** Andrew P. Hoadley, Russell D. Fernald, Beau A. Alward

## Abstract

Steroid hormones play numerous important and diverse roles in the differentiation and development of vertebrate primary and secondary reproductive characteristics. However, the exact role of androgen receptors (ARs)—which bind circulating androgens—in this regulatory pathway is unclear. Teleost fishes further complicate this question by having two paralogs of AR (ARα and ARβ) resulting from a duplication of their ancestral genome. We investigated the functional role of these two ARs on testes growth and development by experimentally eliminating receptor function of one or both paralogs using CRISPR/Cas9 genome edited *Astatotilapia burtoni*, an African cichlid fish. Fish with two or more functional receptor alleles were more likely to be male compared to fish with one or fewer, suggesting that the two paralogs of the receptor may be redundant in regulating early sex determination. In contrast, we found that adult testes size was significantly affected by distinct combinations of mutant and wild-type AR alleles. We present a working model whereby ARβ facilitates testes growth and ARα causes testes regression. This mechanism may contribute to the robust social and physiological plasticity displayed by *A. burtoni* and other social teleost fish.

## Introduction

Androgens play a critical role in the differentiation and maintenance of primary and secondary sexual characteristics in vertebrates (Baroillera et al., 1999). The activity of circulating androgens is mediated by androgen receptors (AR), which, like other steroid hormone receptors, function as ligand-dependent transcription factors (Matsumoto et al., 2008).

Across vertebrates, androgens promote the growth of testes, both during embryonic development and at the onset of reproductive capacity (Alward et al., 2020; Juntti et al., 2010; Walters et al., 2010). In teleost fish, precisely determining the role of androgen signaling in regulating testes growth has been difficult because of a teleost-specific whole genome duplication (TS-WGD), which has resulted in paralogs of several key gene families including the nuclear steroid receptors (Glasauer and Neuhauss, 2014). For example, two isoforms of AR have been described in the Japanese eel (Ikeuchi et al., 1999), Atlantic croaker (Sperry and Thomas, 1999), Western mosquitofish (Ogino et al., 2004), rainbow trout (Takeo and Yamashita, 1999), and the cichlid *Astatotilapia burtoni* (Harbott et al., 2007).

*A. burtoni* are an excellent candidate in which to investigate the role of ARs on testes mass, as they exhibit reproductive plasticity throughout their lives that can be modeled in the laboratory (Fernald, 2012). We recently generated using CRISPR/Cas9 gene editing *A. burtoni* that lack functional ARα, ARβ, or both (Alward et al., 2020). Male *A. burtoni* exist in a social hierarchy, where dominant males possess large testes, bright coloration, and perform aggressive and reproductive behaviors while non-dominant males do not. We found that males lacking ARα possessed larger testes than wild-types, while males lacking ARβ or both receptors possessed smaller testes than wild-types. Questions still remain regarding the role of AR paralogs in the regulation of testes mass in *A. burtoni,* however. For example, it is unclear if the presence or absence of one receptor influences the effects of the other when the other is functionally disabled. Indeed, testes mass has never been analyzed in a species with different combinations of mutations in paralogous ARs. An analysis such as this may yield fundamental insights into the role of paralogous ARs in the control of reproductive plasticity in teleost fish. Here, we addressed this question using *A. burtoni* with multiple combinations of mutated or wild-type AR alleles (Summarized in Fig. 1).

**Figure 1.**
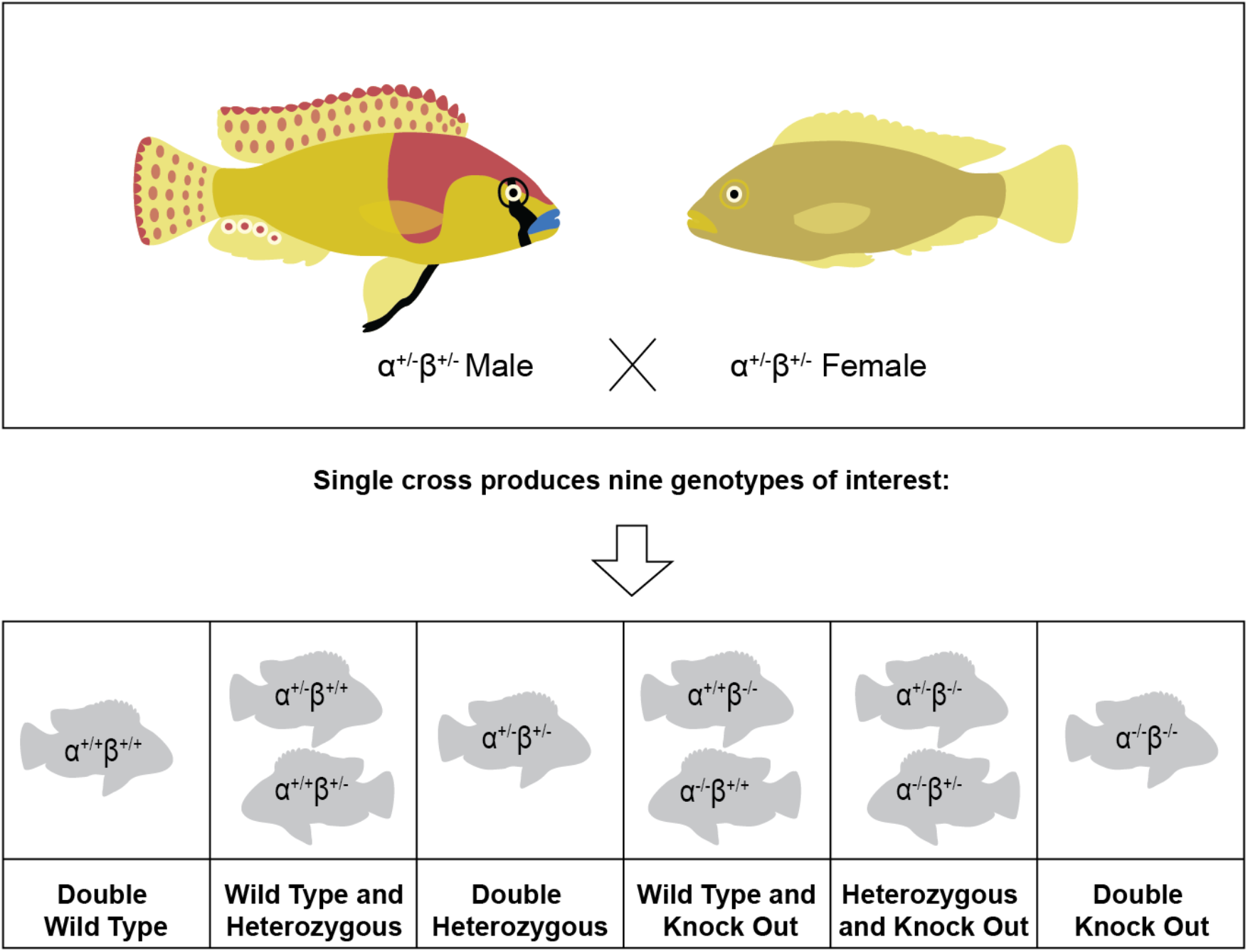
Dihybrid cross of α^+/−^β^+/−^ genotype fish. To generate fish with different combinations of functional and non-functional paralogous AR alleles, two double heterozygous (α^+/−^;β^+/−^) fish were mated, yielding the nine genotype combinations we used in our experiment. Multiple matings were required to produce the 169 individuals that were used in the current study. The overall genotype for fish was written to indicate the allele status of ARα first and ARβ second. For example, for fish heterozygous for both paralogs, their genotype was written as “α^+/−^;β^+/−^”; if they were heterozygous for ARα and wild-type for ARβ, their genotype was written as “α^+/−^;β^+/+^”.

## Materials and Methods

### Experimental Animals

All of the fish used in this experiment were adult *A. burtoni* males descended from a wild caught population from Lake Tanganyika, Africa (Fernald and Hirata, 1977) in accordance with Association for Assessment and Accreditation of Laboratory Animal Care standards. All experimental procedures were approved by the Stanford University Administrative Panel for Laboratory Animal Care (Protocol #9882). The fish were maintained under environmental conditions that mimic their natural equatorial habitat (28 °C; pH 8.0; 12:12 h light/dark cycle with full spectrum illumination; constant aeration). Aquaria contained gravel-covered bottoms with terra cotta pots cut in half to serve as shelters and spawning territories. Fish were fed cichlid pellets and flakes (AquaDine, Healdsburg, CA, USA) each morning.

### Modification of ARα and ARβ using CRISPR/Cas9 gene editing

Mutations in *ARα* or *ARβ* were generated using CRISPR/Cas9 gene editing. The details of the methodology used were described in detail recently in Alward et al (2020). Briefly, the mutant *ARα* allele lacked 50 bps (*ARα^d50^*) within exon 1 upstream of the DNA binding domain (DBD) and Ligand binding domain (LBD) but downstream of the transcription start site, while the mutant *ARβ* allele lacked 5 bps (*ARβ^d5^*) within exon 1 upstream of the DBD and LBD but downstream of the transcription start site. These mutations yielded proteins with premature stop codons and completely lacked the complex tertiary structure seen in the wild-type versions, likely rendering both proteins to be completely non-functional. To generate fish with different combinations of functional and non-functional paralogous AR alleles, we mated male and female fish heterozygous for both receptors, yielding nine genotype combinations (see Fig. 1). Fish were mated multiple separate times to produce a total 169 offspring that were used in the current study. Fish from each mating were housed in separate mixed-sex community tanks (121 L) containing 6-48 fish. For clarity, throughout the manuscript and in the figures, “+” indicates a particular allele is wild-type (also referred to as functional), while a “-” indicates a particular allele is mutated (also referred to as non-functional). Notation of the overall genotype for fish was written to indicate the allele status of *ARα* first and *ARβ* second. For example, for fish heterozygous for both paralogs, their genotype was written as “α^+/−^;β^+/−^”; if they were heterozygous for *ARα* and wild-type for *ARβ*, their genotype was written as “α^+/−^;β^+/+^”.

### Fin Clipping, DNA Extraction, and PCR Amplification

After removing the fish from the tank, they were immediately fin-clipped. Using ethanol-cleaned scissors, a 1-to 2-mm portion of the caudal fin was excised and placed into an individual PCR tube. This was repeated for the rest of the fish run on a given day and the scissors were cleaned thoroughly with ethanol between each fin clipping. To extract DNA, 180 mL of NaOH (50 mM) was added to the sample, which was incubated at 94 °C for 15 min. After incubation, 20 mL of Tris·HCl (pH 8) was added directly into the sample, which was then vortexed and spun down using a minicentrifuge for 5 s. The samples were then placed at −20 °C for at least 15 min before PCR amplification of mutated regions of ARα or ARβ.

For *ARα* we PCR-amplified a 536-bp amplicon spanning the Cas9 target sites with the primers *ARα*FlankF, 5′-CCCAGTGCACTCTAACTCCG-3′ and *ARα*FlankR, 5′-TTTAAGGGTACGACCTCG-GC-3′ and visualized the products on a gel. We performed the same procedure for *ARβ* by PCR amplifying a 642-bp amplicon spanning the Cas9 target site using the primers *ARβ*FlankF, 5′-CCA-TCCCACCTCCAAGAGTC-3′ and *ARβ*FlankR, 5′-GAGGACAGGCCGATGATGAA-3′ and Sanger-sequenced the product with *ARβ*FlankF (Lone Star Labs, Houston, TX).

### Measuring testes mass

Fish were removed from their aquaria and their standard length and body mass were recorded. Fish were then killed by cervical transection and the testes were removed and weighed. Testes mass was standardized to body mass by calculating the gonadosomatic index (GSI=[testes mass/body mass]*100).

### Statistical Analysis

Statistical analyses were performed using GraphPad Prism version 9.0.1 (Mac OS X, GraphPad Software, San Diego, California USA) or RStudio (Version 1.2.5019). The sample was compared to the expected genotype distribution based on a dihybrid cross using Chi-squared tests with simulated p values due to the small sizes of some groups. The effects of genotype on sex were analyzed using a Chi-square test followed by Fisher’s Exact Tests corrected for false discovery rate (fdr) for paired comparisons. GSI values were transformed using the square root of these values to meet the assumptions for a parametric One-Way ANOVA. Kruskal-Wallis ANOVAs were used to assess SL and BM values because they did not meet the equality of variance assumption regardless of attempts to transform the values (Brown-Forsythe test, p<0.05). Following a significant main effect for an ANOVA, Tukey’s (parametric) or Dunn’s (Kruskal-Wallis) tests were used for pairwise comparisons. Effects were considered significant at p≤0.05. Data and code used are available at https://github.com/AlwardLab.

## Results

### Sex is influenced by AR paralogs

Distribution of genotypes did not significantly differ from Mendelian ratios expected by a dihybrid cross (χ^2^=9.127, df=NA, p=0.323). Qualitative analysis of the effects of genotype on sex revealed findings relevant for subsequent analyses on GSI. For example, only two fish (out of 7) possessing no functional AR alleles were male (Fig. 2A). We do not believe the low sample size of males in this group poses issues for our later analyses. Indeed, our central focus here was on the role of combinations of different functional AR alleles on testes mass and we have recently shown the effects of possessing no functional AR alleles on testes mass (Alward et al., 2020).

**Figure 2.**
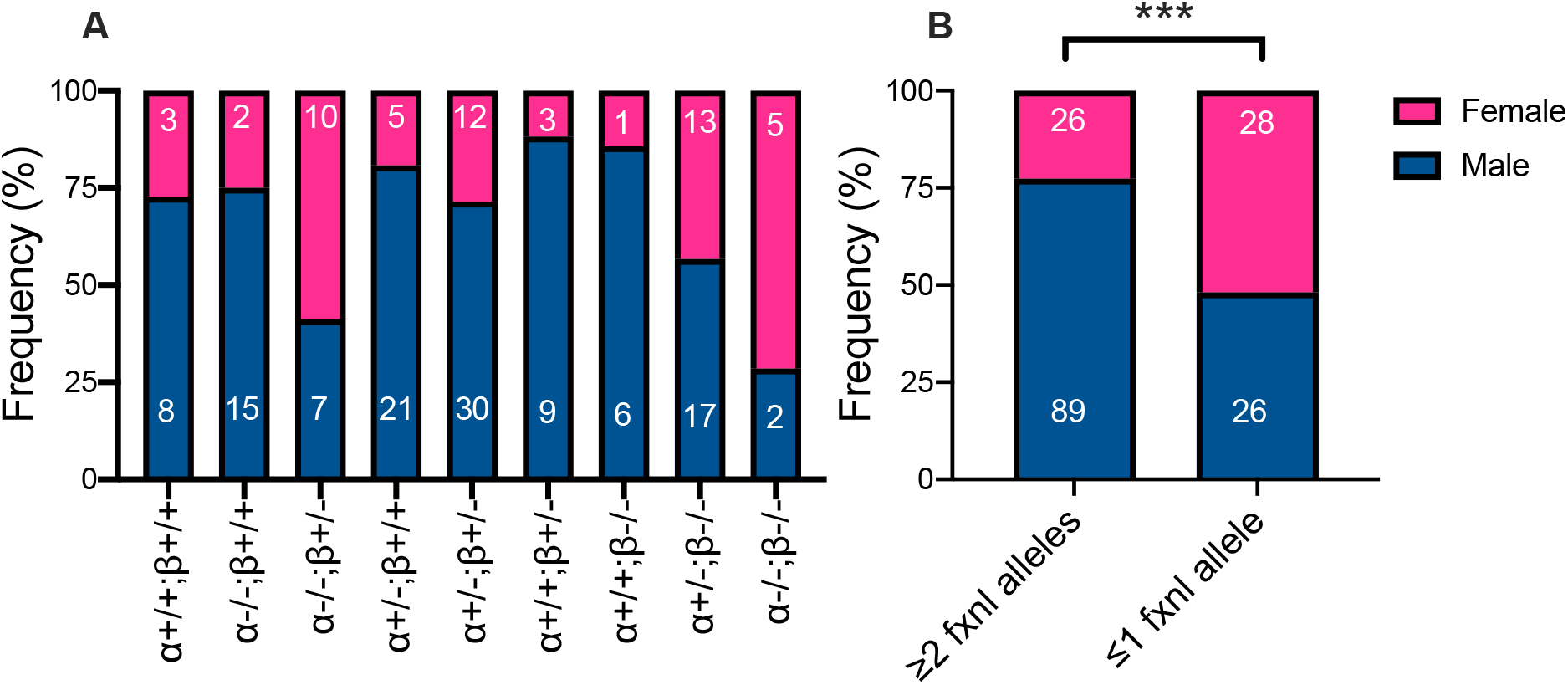
Effects of AR mutations on sex. (A) Frequency of males and females across genotypes. (B) Frequency of males and females after grouping fish into possessing two or more functional AR alleles or one or zero functional alleles. Numbers in white font over the male or female portion of the bars are actual numbers of fish. fxnl=functional. \*\*\**P* < 0.001 (Fisher’s Exact Test).

A Chi-square test yielded a significant effect of genotype on sex (Fig 2A; χ^2^=19.17, df=8, p=0.01). Post-hoc tests were unable to identify which differences drove the significant omnibus Chi-square test, likely because the fdr-corrected critical p-value was so low due to the large number of paired comparisons conducted. Upon visual inspection of Figure 1A it appeared that fish with two or more functional AR alleles were more likely to be male compared to fish with one or fewer functional AR alleles, which could explain what was driving the significant omnibus Chi-square test. Therefore, we collapsed across these categories and compared the two groups using a Fisher’s Exact Test. We found that fish with two or more functional AR alleles were significantly more likely to be male than fish with one or fewer functional AR alleles (Fisher’s, p=0.0003), suggesting two functional AR alleles, regardless of which paralog, are sufficient for increasing the likelihood of being male in *A. burtoni*.

### GSI is significantly affected by distinct mutant and wild-type AR allele combinations

We observed a significant effect of AR paralog mutation on GSI (*F*_8,106_= 22.20; *P*<1*10^-15^). All groups with fish with zero functional *ARβ* alleles—regardless of the mutational state of *ARα*—had significantly smaller GSI compared to all other groups except for *ARα^+/+^;ARβ^+/−^* males and did not differ from one another (Fig. 3). *ARα^+/+^;ARβ^+/+^* fish did not significantly differ from fish with any combination of homozygous mutant *ARα*, heterozygous *ARα* or *ARβ*, or wild-type *ARβ*. In addition to their noted differences from all *ARβ^−/−^* males, *ARα^−/−^;ARβ^+/+^* males had significantly larger GSI compared to *ARα^+/+^;ARβ^+/−^* and *ARα^+/−^;ARβ^+/−^*. *ARα^−/−^;ARβ^+/−^* and *ARα^+/−^;ARβ^+/+^* males only differed significantly form the *ARβ^−/−^* males. Finally, there were no effects of AR mutation on BM (*KW*: 10.46, *P*=0.23) or SL (*KW*: 10.46, *P*=0.23) (Fig. 4A-B). This pattern of results highlights a potentially complex relationship between the presence of functional AR paralogs and the control of testes mass.

**Figure 3.**
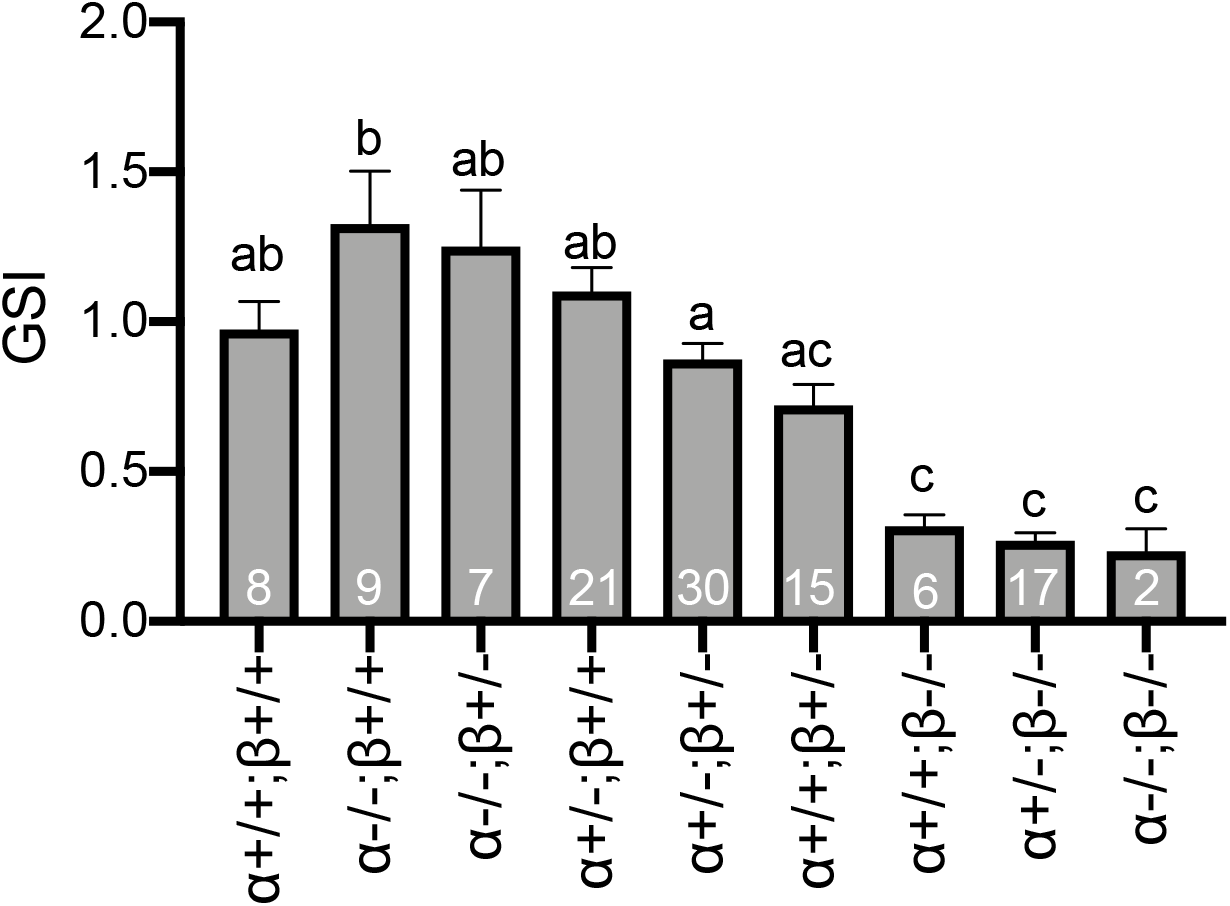
Effects of AR mutations on GSI. There was a significant effect of genotype on GSI in males. Groups with the same letters written above them are not significantly different from one another. Numbers in white on each bar indicate number of fish with that genotype. Bars are mean ± SEM.

**Figure 4.**
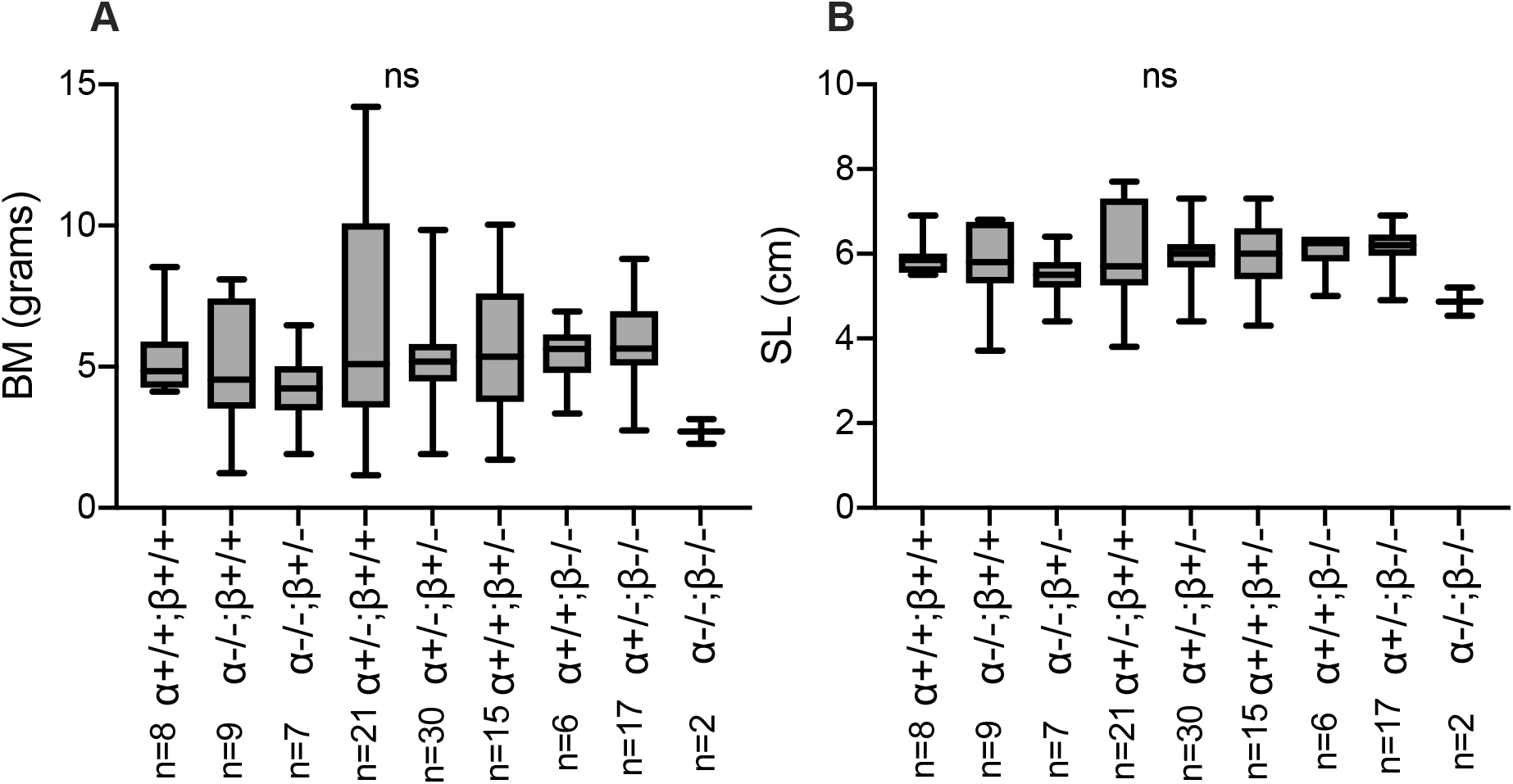
No significant effects of AR mutations on body size measures. There was no effect of genotype on (A) BM or (B) SL. Bars represent median with minimum, maximum, first, and third quartiles. BM=body mass; SL=standard length. Numbers below each genotype indicate sample size. ns=non-significant.

**Figure 5.**
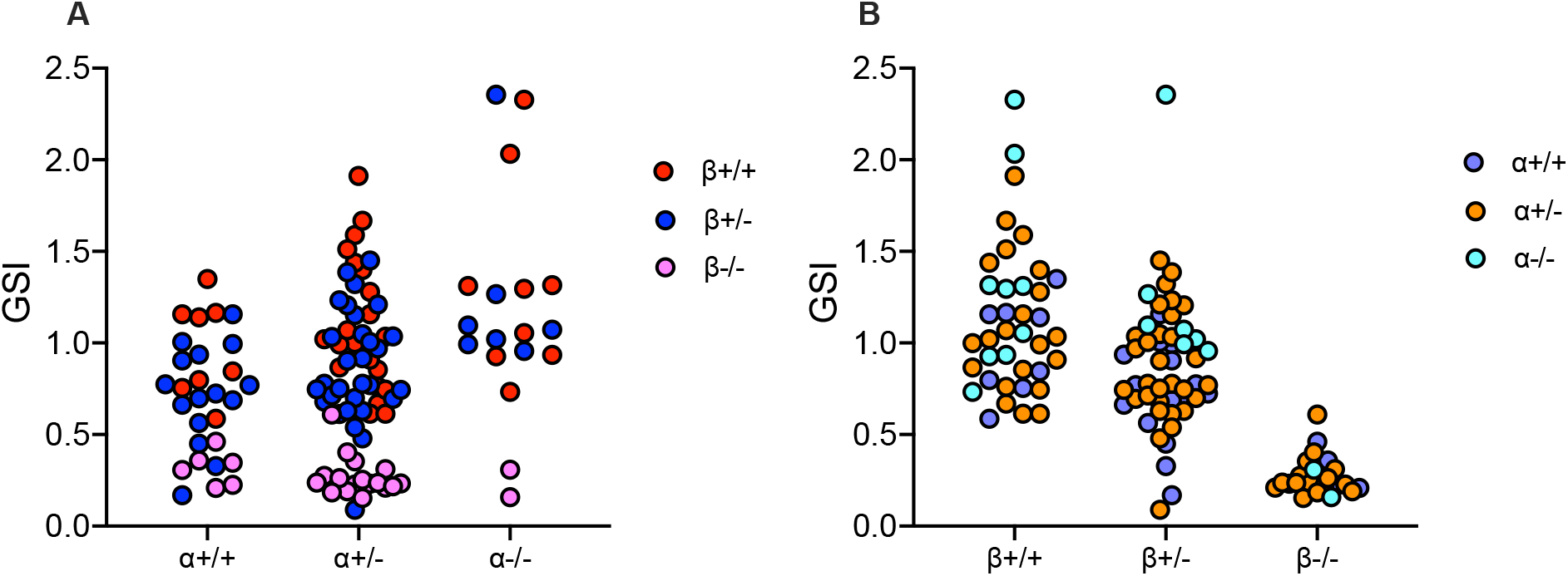
ARβ is permissive for the effects of ARα mutations on GSI. GSI values were potted for (A) *ARα* genotypes as a function of *ARβ* and (B) the same was done for *ARβ*. As shown in (A), no matter the state of *ARα*, if fish did not possess any functional *ARβ*, GSI remained low. (B) The same was not true for *ARβ*.

These results suggest the mutational state of *ARβ* determines the extent to which the mutational state of *ARα* affects GSI size. To gain a clearer picture of this relationship, we plotted the effects of the mutational state of *ARα* as a function of *ARβ* and vice-versa (Fig. 4). These data show clearly that regardless of *ARα* mutational state, if fish possess zero functional *ARβ* there they have very small testes (Fig. 4A). On the other hand, the fewer functional *ARβ* alleles a fish has, the smaller testes they possess, regardless of *ARα* mutational state (Fig. 4B). Thus, functional *ARβ* is required for the enhancement in GSI induced by *ARα* mutation, but *ARα* is not required for the reduction in GSI cause by *ARβ* mutation.

## Discussion

We have shown that AR paralogs in *A. burtoni* are involved in sex determination and control testes growth. Specifically, we show evidence that either AR paralog may play a role in biasing fish toward male: regardless of which paralog was mutated, possessing two functional versions of either one was sufficient for this bias. Moreover, we provide further confirmation that ARα is required for reducing testes growth and ARβ is required for enhancing testes growth (Alward et al., 2020). Our results have important implications for understanding the androgenic control of sex and testes growth in *A. burtoni* and other vertebrates.

We have shown recently that ARα and ARβ exert non-redundant control of coloration, behavior, and testes mass (Alward et al., 2020). The current results expand on this idea, providing more precise evidence of subfunctionalization of the ARs (Glasauer and Neuhauss, 2014; Ogino et al., 2016). In our case, ARα reduces testes growth and ARβ promotes testes growth. Moreover, our results suggest that the control of testes mass by ARα likely involves a reduction in ARβ function. Future experiments investigating the potential interactions between ARα and ARβ will be important to elucidate these mechanisms.

The control of sex in *A. burtoni* is hypothesized to be highly polygenic (Roberts et al., 2016). Our data suggest that AR paralogs may be involved. These results are in line with the findings across several fish species that treatment of larval fish with androgens can significantly bias them towards developing as male (Baker et al., 1988; Blazquez et al., 2001; Feist et al., 1995; Gale et al., 1999; Kavumpurath and Pandian, 1994; Marjani et al., 2009; Örn et al., 2003; Pandian and Sheela, 1995). The current findings are especially relevant to those studying the effects of early androgen exposure on sexual development in teleost fish that also have two AR paralogs. Indeed, our AR mutant *A. burtoni* are an excellent model in which to use androgen exposure methods to pinpoint precisely which AR is involved in different processes underlying sexual differentiation and maturation.

We show that the ARα and ARβ appear to serve redundant roles in the control of sexual differentiation but non-redundant roles in the control of adult testes growth, highlighting the complex modular roles of AR paralogs in *A. burtoni* in the control of physiology and behavior (Alward et al., 2020). We have developed working model of the control of testes mass by ARs in *A. burtoni* that is summarized in Figure 6. The hypotheses presented in this model present clear avenues for future research.

**Figure 6.**
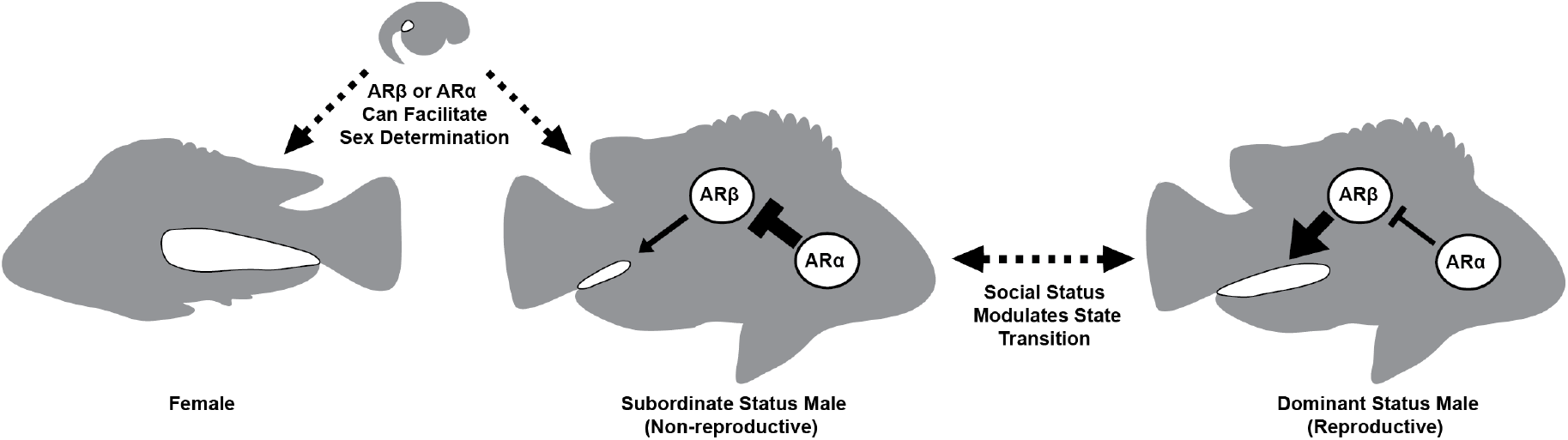
Working model of testes size regulation by AR paralogs. ARα and ARβ appear to serve redundant roles in the control of sexual differentiation but complementary roles in the control of adult testes growth. Two functional alleles of AR, regardless of which paralog, may be sufficient to skew sexual differentiation towards males. In contrast, ARα and ARβ may have opposing effects on adult male testes size. For example, in our model subordinate males are predicted to have low ARβ activity due to inhibition by ARα, which leads to small testes. In dominant males, ARβ activity is high due to reduced inhibition by ARα. This novel mechanism may contribute to the striking abilities of A. burtoni males to transition between reproductive and non-reproductive states as a response to social opportunity.

## Conclusion

The diversity among teleost fish has garnered the attention of researchers from numerous fields, including genetics and evolution. It has been proposed recently that the incredible adaptive radiation among a particularly teleost clade, the African cichlid fish, is partly due to the presence of duplicated genomic regions from the TS-WGD (Brawand et al., 2014). Some have suggested that the duplication of AR paralogs specifically may have been especially important in driving the diversity in social systems among cichlids (Alward et al., 2020; Douard et al., 2008; Lorin et al., 2015). Our experiments in AR mutant *A. burtoni* support these ideas, showing clearly the complex, non-redundant roles played by AR paralogs in the control of suites of traits related to reproduction that may contribute to the remarkable social plasticity displayed by this species.

## Acknowlegements

We thank Vibhav Laud (Stanford University) for assistance dissecting fish. We also thank Humayd Mirza and Rohail Siddiqi (University of Houston) for assistance with PCRs and genotyping. Lastly, we thank Mariana Lopez and Lillian Jackson (Unviersity of Houston) for providing feedback on an earlier version of this manuscript.

